# Chromatin assembly at replication forks enables differentiation of human iPSCs

**DOI:** 10.64898/2026.01.20.700317

**Authors:** Jane Wright, Hao Jiang, Susanne Bandau, Simone Di Sanzo, Lindsay Davidson, Moritz Volker Albert, Angus Lamond, Constance Alabert

**Affiliations:** Division of Genome Integrity, School of Life Sciences, University of Dundee, UK; Division of Molecular, Cell C Developmental Biology, School of Life Sciences, University of Dundee, UK; MOLEQLAR Analytics GmbH, München, Germany; Human Pluripotent Stem Cell Facility, School of Life Sciences, University of Dundee, UK

## Abstract

DNA replication is both productive and disruptive, synthesising new DNA while dismantling chromatin to allow fork progression. Because of its disruptive nature, DNA replication has been suggested to enable chromatin changes facilitating cell fate transitions. However, the role of DNA replication in cell fate transitions remains incompletely understood. Here we show that upon differentiation of human induced Pluripotent Stem Cells (iPSCs) into presomitic mesoderm, DNA replication allows hundreds of proteins, including transcription factors, histone chaperones, and histone methyltransferases, to access chromatin. Moreover, H3K9 methylation is enhanced upon differentiation through a DNA replication-coupled process. Most of the proteins recruited to (or evicted from) nascent chromatin during differentiation are already present in iPSCs, suggesting they may play a role in the early response to differentiation. Among these, we identified ERH and SETDB1 as essential for mesoderm differentiation. Our work reveals a new paradigm to explore: how differentiation signals exploit the replisome interactome to enable rapid chromatin changes.

## INTRODUCTION

DNA replication is a fundamental biological process essential for the faithful transmission of genetic and epigenetic information [1–3]. This process involves the core replisome and a network of replication-associated proteins, chromatin modifiers, and signaling pathways that are frequently disrupted in diseases such as cancer and neurodevelopmental disorders [4–10]. Replisome progression provokes genome-wide disassembly of chromatin ahead of replication forks, thereby offering an opportunity for epigenetic change [1–3, 11]. Moreover, recent evidence supports a causative link between DNA replication and cell fate transitions [12–19]. The underlying mechanisms are poorly understood, and a variety of possibilities have been proposed, ranging from the disruption of pre-existing chromatin determinants [13, 17–19] to the opportunity for assembling distinct chromatin states *de novo* [12, 14–16, 20]. Consistent with the latter possibility, we and others showed that DNA replication allows hundreds of transcription factors and chromatin effectors, such as histone methyltransferases, to bind transiently to the genome in human cell lines [21–24]. However, for many of these proteins, it remains unknown whether their recruitment to areas of nascent chromatin facilitates changes that are functionally important for cell fate transitions.

To study the role of DNA replication during cell fate transitions, we have established the ‘iPOND’ protocol (isolation of Proteins On Nascent DNA [25]) in human induced pluripotent stem cells (iPSCs), both in the pluripotent cell state and upon differentiation. When human iPSCs are exposed to signals that induce differentiation into presomitic mesoderm (PSM) [26, 27], we observe a profound and rapid rewiring of the proteins that are recruited around the human replisome. These proteins are a mixture of transcription factors, chromatin regulators, and some proteins with no previously known function in cell fate transitions. Importantly, a large proportion of these proteins are already expressed in iPSCs and thus are potential “first responders” to developmental cues. Moreover, a change in H3K9 histone methyltransferases interacting with the replisome is concurrent with a change in the restoration of di- and trimethylation on histone H3 on lysine 9 (H3K9me2, H3K9me3) behind replisomes. Finally, we show that PSM differentiation progression requires transit through S-phase, DNA replication coupled chromatin assembly, as well as two of the first responders identified in this study, SETDB1 and ERH. Altogether these findings suggest differentiation signals rapidly re-wire newly replicated chromatin and shed light on the mechanisms enabling the coordination between DNA replication and differentiation.

## RESULTS

### Rapid rewiring of the replisome interactome upon differentiation

To explore how DNA replication-coupled mechanisms can contribute to cell differentiation, we first identified proteins that differentially associate with replicating chromatin during differentiation. To do so we compared the composition of newly replicated chromatin in human iPSCs versus PSM induction (Figure 1A). Differentiation was induced by incubating iPSCs in CL medium containing a GSK3 Inhibitor/WNT mimetic (CHIR99021) and a BMP inhibitor (LDN193189) for up to 72 hours according to established protocols [26, 27]. As expected, most iPSCs are in S phase, and by 72 hours of PSM differentiation the cell population start to accumulate in G1 phase of the cell cycle (Figure S1A, S1B and S1C). After 24 hours in CL medium, cells acquired a neuromesodermal progenitor (NMP) [28–30], primitive streak-like fate, characterized by expression of TBXT (Brachyury), and after 48 hours a PSM fate, marked by TBX6 expression in most cells (Figure 1B, S1D-F). From 48 hours most cells showed low expression of pluripotency markers NANOG and SOX2 (Figure 1B, S1D-F). Expression of differentiation markers therefore occurs before complete silencing of pluripotency markers takes place [31, 32], and within 48 hours iPSCs exit pluripotency and become PSM.

**Figure 1:**
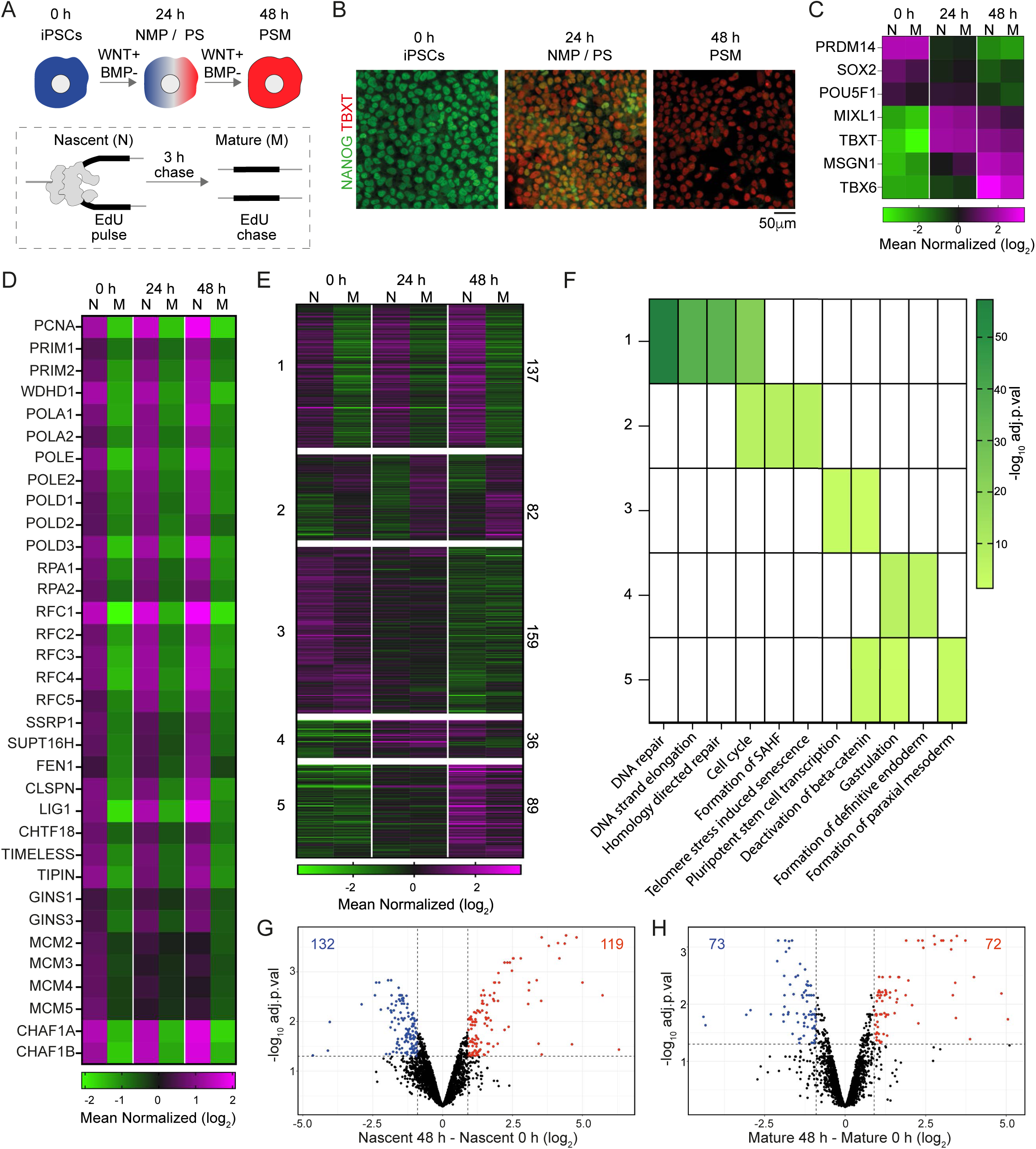
Rapid rewiring of the replisome interactome upon differentiation. A. Schematic for the presomitic mesoderm (PSM) differentiation pathway and the Isolating proteins on nascent DNA (iPOND) assay. Induced pluripotent stem cells (iPS) and cells differentiated along the PSM pathway for 24 or 48 hours are treated with the nucleotide analog EdU which incorporates into nascent DNA. Half of cells are harvested immediately to identify proteins associated with nascent (N) chromatin or in the other half, EdU is washed out, cells chased with thymidine and then isolated 3 hours later to identify proteins associated with Mature (M) chromatin. B. Representative immunofluorescence images of a pluripotency marker (NANOG) and an early mesodermal marker (TBXT/Brachyury) at 0, 24, and 48 hours of PSM differentiation. C. Heatmaps showing differential binding of cell type specific transcription factors to nascent and mature chromatin during PSM differentiation. Scale is mean normalized log_2_ (reporter intensities). D. Heatmaps of known replisome proteins across all iPOND samples. Scale is mean normalized log_2_ (reporter intensities). E. Clustering (K-means) of any significantly changing protein among iPSCs, 24 hours, and 48 hours PSM nascent and mature chromatin. Significantly changing proteins were defined to be all proteins with an adjusted p value <0.05 and a log_2_ fold change of at least 0.9 in any given pairwise contrast (n=503 proteins). Scale is mean normalized log_2_ reporter intensities. The number of proteins detected in each cluster is indicted on the right. F. Top Reactome pathway terms (REACC) detected for each cluster from panel E and their representation in all clusters. Significance is shown in increasing green colour reflecting minus log_10_ adjusted p value with 1.3 the minimum significant value and non-significance shown in white. G. Volcano plots generated from the iPOND-TMT-MS data for comparison of Nascent chromatin. Limma *t* test revealed significantly changing proteins with a log_2_ fold > +/- 0.9 and p.adj <0.05 are shown in blue (iPSCs) or red (48 hours PSM). H. Same as G but for comparison of Mature chromatin between iPSCs (blue) and 48 hours PSM cells (red). See also Figure S1 and Table S1

To isolate newly replicated chromatin, the iPOND protocol was established in human iPSCs [25]. In brief, cells were incubated for 20 minutes with the nucleotide analog EdU which incorporates into replicating nascent DNA [25, 33]. Half of the cells were then processed immediately to isolate EdU-labelled nascent chromatin (N), while the other half were incubated with thymidine for an additional 3 hours before the cells were collected (Figure 1A). The thymidine chase blocks EdU incorporation and allows isolation of EdU-labelled matured chromatin (M), allowing to discriminate proteins enriched on chromatin in general (M) from proteins enriched on nascent chromatin (N). Proteins associated with replicated chromatin were identified by immunoblot (Figure S1G), and by Tandem Mass Tag mass spectrometry (TMT-MS) on three biological iPOND replicates (Figure 1C-E, Figure S1H, Table S1). In total we identified 2956 proteins that were detected above the background of non-EdU exposed cells (see methods). Pluripotency markers were enriched on replicated chromatin in iPSCs and differentiation markers enriched in PSM cells (Figure 1C). Moreover, core replisome components were enriched on nascent chromatin compared to mature chromatin in iPSCs, PSM-24 hours, and PSM-48 hours indicating that newly replicated chromatin was successfully isolated (Figures 1D and S1G).

503 out of 2956 proteins detected are differentially associated with replicated chromatin during 48 hours of PSM differentiation (17% of the proteins identified; see methods). Clustering the 503 proteins revealed five distinct clusters (Figure 1E, Table S1). Cluster 1 contains proteins known to be involved in DNA replication and repair, and Cluster 2 contains proteins involved in Cell cycle and Senescence associated heterochromatin formation such as histone H1 [21] (Figure 1F, Table S1). Proteins from clusters 1 and 2 (44% of the 503) show no change of association patterns to replicated chromatin upon differentiation, either consistently increased on nascent chromatin (Cluster 1) or consistently increased on mature chromatin (Cluster 2), revealing that these processes are not affected by differentiation. In contrast, 56% of the 503 proteins (Cluster 3, 4, and 5) show differential enrichment behind replisomes upon differentiation (Figure 1E). Proteins in Cluster 3 (159 proteins) are enriched on iPSCs chromatin and include proteins involved in pluripotent stem cell transcription (Figure 1F). Conversely, proteins in cluster 4 and 5 are enriched on PSM chromatin, with those that peak at 24 hours (36 proteins) enriched for the REACTOME terms, Gastrulation and Endoderm, and those peaking at 48 hours (89 proteins) associated with the terms Gastrulation and Formation of paraxial mesoderm (Figure 1F). Altogether this analysis reveals that upon differentiation, the composition of proteins associated with replicated chromatin changes.

Next, we performed a direct comparison of the composition of nascent chromatin upon differentiation. Strikingly, at 48 hours, 119 proteins are newly recruited to nascent chromatin and 132 are lost (Figure 1G). The composition of mature chromatin is less affected: 72 proteins are gained and 73 are lost in PSM cells compared to iPSCs (Figure 1H). Moreover, from the 119 proteins recruited to nascent chromatin upon differentiation, 52 are also enriched on mature chromatin whereas 67 are transiently enriched on nascent chromatin (Figure S1I). Therefore, as iPSCs undergo differentiation, there is a rapid change in composition of replicated chromatin with most changes taking place on nascent chromatin.

### Distinct groups of transcription factors (TFs), epigenetic modifiers, and histone chaperones are recruited to newly replicated chromatin upon differentiation

Almost one quarter of the 503 significantly changing iPOND proteins are classified as TFs according to TF Database 4.0 (121/503; [34]) (Figure 2A). Notably the PSM-enriched factors (cluster 4 and 5, Figure 1E) contain a large proportion of TFs (44%), including mesoderm lineage specific TFs MSGN1 and TBX6. Importantly, most TFs, including pioneer TFs, have increased association with nascent chromatin regardless of their DNA binding domain (Figure 2B, S2A) [34–36]. Previous work in other contexts suggested that DNA replication may provide a "window of opportunity" for establishing new TF binding patterns in daughter cells [37]. This work reveals that the increased association of TFs with nascent chromatin must rely on additional features and mechanisms other than pioneer status or specific DNA binding domain.

**Figure 2:**
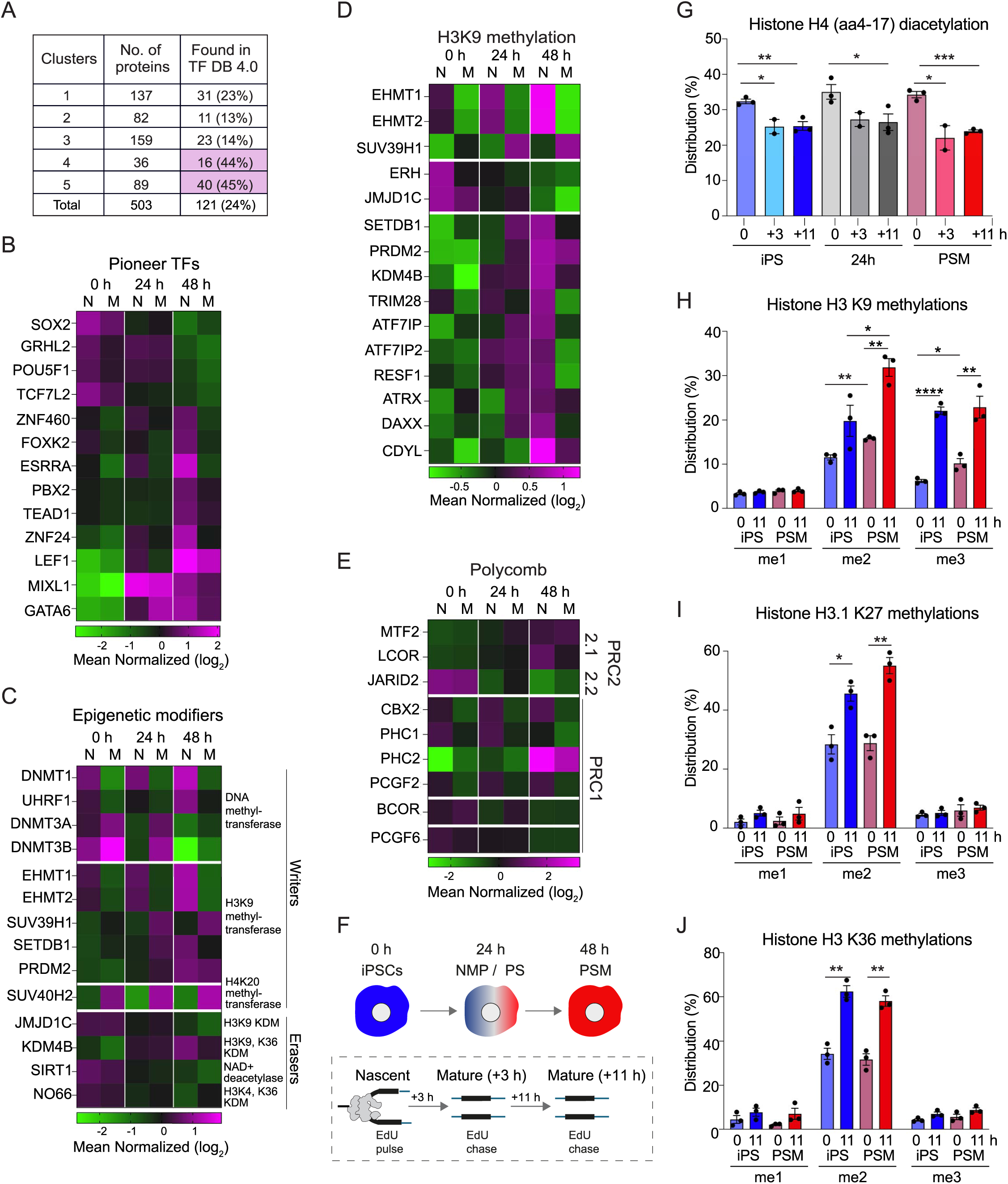
Restoration of heterochromatin behind replisomes changes upon differentiation. A. Using the TFDB4.0 database [34] 121 transcription factors (TFs) are identified within the 503 significantly changing proteins. These TFs are grouped according to the 5 clusters shown in Figure 1E to see the prevalence of TFs per cluster. B. Heatmaps showing differential binding of Pioneer Transcription factors to nascent and mature chromatin during PSM differentiation. Scale is mean normalized log_2_ (reporter intensities). Protein list is taken from [35, 36]. For the heatmaps in B-E only proteins detected within the list of 503 significantly changing proteins in the iPOND-TMT experiments are shown. C. Same as B. but for the Epigenetic writers and erasers of histones H2A, H2B, H3 and H4. D. Heatmaps showing differential binding of proteins involved in H3K9 methylation to nascent and mature chromatin during PSM differentiation. Scale is mean normalized log_2_ (reporter intensities). E. Same as for D but focusing on polycomb proteins that are part of PRC1 or PRC2 complexes. F. Experimental design to detect histone modifications in nascent and mature chromatin in iPSCs and cells differentiating along the PSM pathway. Newly replicated DNA was labelled with EdU and isolated by iPOND and enriched histone modifications subsequently identified by mass spectrometry. G. Enrichment of di-acetylated histone H4 on nascent chromatin samples validates that they were undergoing replication. The level of each histone modification identified on a residue is expressed as a percentage of all the possible modified states detected for this residue [51]. Diacetylation level on the H4 amino acid 4–17 (aa4–17) peptide in nascent (0 h) and mature chromatin (+3 and +11 h). H4 aa4-17 can be acetylated on K5, K8, K12, and K16. Level of diacetylation on H4 aa4-17 is shown as a percentage of all the aa4-17 peptides detected. H. Dynamics of H3K9 methylation on replicated chromatin in iPS and PSM 48-hour cells. Here, monomethylation (me1), dimethylation (me2) and trimethylation (me3) of H3K9 in nascent (0 h) and mature chromatin (+11 h) are shown as a percentage of all the H3 aa9-14 peptides detected. I. Same as H, but for H3.1 K27 methylations. J. Same as H, but for H3.1 K36 methylations. See also Figure S2 and Table S2

The histone chaperones known to play a role in histone delivery to newly replicated DNA [38–40] (MCM2, SUPT16H, SSRP1, CHAF1A, CHAF1B, TONSL and UBN2), were consistently associated with nascent chromatin in iPSCs and at 24 or 48 hours after differentiation, revealing that this process does not change upon differentiation (Figure S2B). However, the H3.3 histone chaperones DAXX, ATRX, and the HIRA complex showed enrichment on nascent chromatin at 48 hours of PSM induction. Therefore, upon differentiation, distinct histone chaperones may deliver histones to newly replicated DNA.

Several epigenetic modifiers were also identified within the 503 significantly changing proteins. Initially only the catalytic subunits of epigenetic writers and erasers were searched and most of those identified are associated with gene repression (Figure 2C). Broader analysis of these repressive epigenetic modifiers, including noncatalytic subunits, revealed fifteen proteins linked to H3K9 methylation associate differentially with replicating chromatin during differentiation. Consistent with previous studies [21–23], H3K9me1 histone methyltransferases EHMT1 and EHMT2 are strongly enriched on nascent chromatin in iPSCs and PSM cells, whereas SUV39H1 consistently associates with mature chromatin (Figure 2D). However, ten of these fifteen proteins show PSM-48-hour nascent chromatin enrichment, including the histone methyltransferases SETDB1 and PRDM2, while two proteins, Enhancer of Rudimentary Homolog (ERH): a protein recently shown to be required for H3K9me3 maintenance [41, 42], and the demethylase, JMJD1C are both enriched on iPSCs nascent chromatin. This differential association of H3K9 writers and erasers suggests that after replication fork passage, H3K9 methylation restoration and constitutive heterochromatin is established differently in iPSCs and PSM cells. Another group of proteins with binding to replicated chromatin changing upon differentiation is the polycomb complex PRC2 that is implicated in cellular identity, including stem cell maintenance and differentiation [43–46]. Upon PSM induction, there is a switch from PRC2.2 to PRC2.1 subunits, with the PRC2.2 subunit JARID2 enriched on iPSCs nascent chromatin and PRC2.1 subunits (MTF2; LCOR (PALI1)) on PSM-48-hour nascent chromatin (Figure 2E). For PRC1, the CBX2 and PCGF2 subunits consistently associate with nascent chromatin in every cell type, while PHC1 is enriched on iPSCs nascent chromatin and PHC2 on 48-hour nascent chromatin. Interestingly, PHC1 is also thought to activate NANOG transcription and PHC2 is implicated in clustering and condensate formation [47–49]. Together these analyses strongly suggest that the mechanism of establishing constitutive and facultative heterochromatin on newly replicated chromatin changes upon differentiation and potentially in a replication coupled manner.

### Differential recruitment of epigenetic writers on newly replicated chromatin correlates with a change in replication-coupled histone modification restoration

The recruitment of distinct epigenetic writers and histone chaperones on newly replicated chromatin upon PSM differentiation suggests that the establishment of certain histone modifications, such as H3K9me2/3, could be altered upon differentiation in a replication coupled manner. To test this hypothesis, we directly tracked the re-establishment of histone modifications on histones H3 and H4, in iPSCs and PSM-differentiating cells. To do so, we repeated the iPOND approach on iPSCs, PSM 24-hour, and PSM 48-hour cells (Figure 2F). An extra mature time point (11 hours) was included as histone modifications takes several hours to be restored [50–52]. Moreover, histones from whole cell pellets were analyzed in parallel to monitor overall changes in level of histones modifications upon differentiation (Figure S2C). We observed higher levels of diacetylated H4 on nascent chromatin (33%) compared to mature chromatin (25%) and to histones from whole cell pellets (∼14%) (Figure 2G, Table S2), supporting that nascent chromatin was successfully captured [53].

From 35 histone methylations and acetylations analysed, 16 were significantly changing upon PSM induction in whole cell extracts (Figure S2D, S2E, Table S2), of which only seven could be modified through a replication-coupled process (iPOND): 3 histone methylations (H3K4me1, H3K9me2 and H3K9me3) and 4 histone acetylations (H3K9ac, H3K18ac, H3.1K27ac and H4 aa4-17 3ac). Upon PSM induction, H3 acetylation on lysine 9, 18 and 27 decreased on nascent chromatin (Figure S2F) revealing that these acetylation’s could be lost in a replication-coupled manner. In contrast, H3K9me2 and H3K9me3 levels increased on nascent chromatin upon differentiation, as well as H3K9me2 on mature chromatin (Figure 2H), suggesting that upon differentiation these modifications are more rapidly imposed in a replication coupled manner. Finally, although H3K27me1, me2 and me3, as well as H3K36me2 and me3 levels change upon differentiation in whole cell extracts (Figure S2D, S2E), their restoration dynamic behind the fork remained similar in iPSCs and PSM cells (Figure 2I, 2J). H3K27me2 showed a slightly enhanced accumulation on mature chromatin in PSM cells compared to iPSCs (Figure 2I), yet these changes were not significant. Altogether these results revealed that differentiation signals provoke the rapid recruitment of a distinct set of H3K9me writers to newly replicated chromatin and enhance H3K9me2 and H3K9me3 restoration behind the replisomes.

### Upon differentiation, lineage-specific TFs are both expressed and recruited to nascent chromatin, whereas most epigenetic regulators are simply recruited

We next investigated how proteins may access newly replicated chromatin upon differentiation. Single cell transcriptomics have shown that upon PSM induction many genes are rapidly transcribed [27]. Therefore, we wondered to what extent new protein synthesis may drive changes in the replisome interactome during differentiation. To look at this, we monitored changes in overall abundance of proteins in whole cell extracts (WCE) upon differentiation (Figure 3A), and contrasted these with changes in replicated chromatin (iPOND) composition as cells undergo differentiation (Figure 1). 6955 proteins were identified, and using the same criteria as for replicated chromatin (see methods), we found that 881 proteins significantly changed in abundance within 48 hours of PSM differentiation (Table S3, Figure S3A). Comparing only iPSCs and PSM-48-hour WCE, 825 proteins change significantly, representing 12% of all the proteins identified (Figure 3A), including pluripotency and differentiation markers as expected (Figure 3B) [27]. Strikingly, only a third of the proteins whose association to replicated chromatin changes upon differentiation also change in WCE abundance (182 out of 503, Figure 3C). However, most changes observed on nascent chromatin (321 out of 503) during differentiation are not coupled to a change in WCE abundance. Therefore, the differential association of 2/3 of proteins to replicated chromatin is unlikely driven by changes in expression.

**Figure 3:**
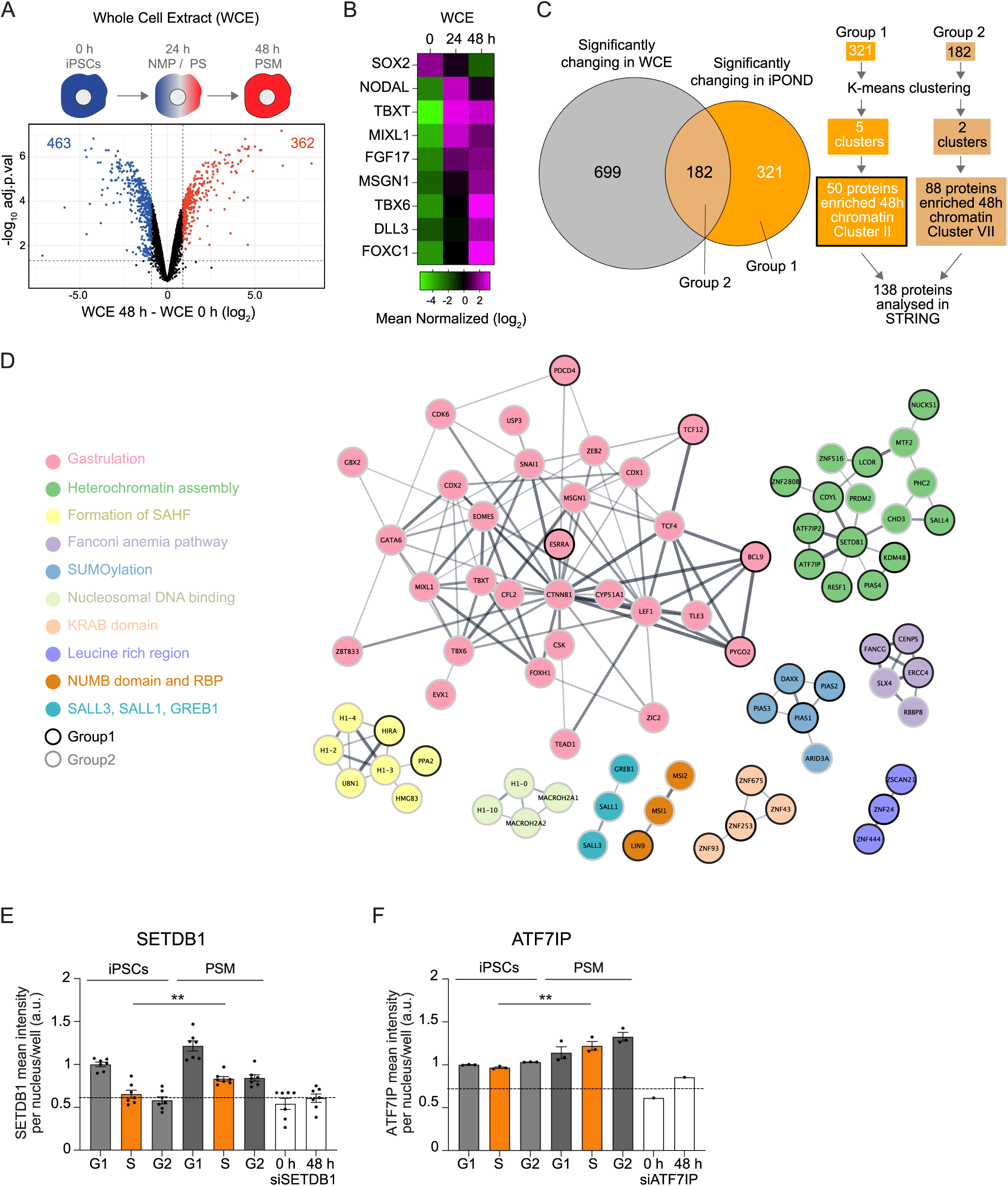
Majority of replisome-associated proteins associate with newly replicated chromatin during differentiation without an increase in protein abundance. A. Schematic for the presomitic mesoderm (PSM) differentiation pathway and a volcano plot generated from the iPOND-TMT-MS data for comparison of proteins detected in Whole Cell Extract (WCE). Limma *t* test revealed significantly changing proteins with a log_2_ fold > +/- 0.9 and adjusted p value <0.05 are shown in blue (iPSCs) or red (48 hours PSM). B. Heatmaps showing differential binding of selected markers of paraxial mesoderm differentiation in the WCE samples at 0, 24 and 48 hours of PSM differentiation. Scale is mean normalized log_2_ (reporter intensities). Protein list is taken from [27]. C. Left: Venn diagram between significantly changing proteins from the WCE (grey) and the iPOND-TMT experiments (orange). Right: a schematic pipeline of the clustering, GO, and STRING/CYTOSAPE analysis for the 182 proteins that are significantly changing in both the WCE and the iPOND experiments as well as the 321 proteins who change significantly in only the iPOND experiments. See also Figure S3 A-C. D. STRING interaction network of the 138 (88 plus 50 proteins indicated in 3C) proteins enriched on PSM chromatin. Only high confidence interactions were considered. Network was imported from STRING into Cytoscape. The nodes that are outlined in black are the proteins that do not change significantly in WCE but do significantly change in the iPOND data (Group 1). The nodes outlined in grey are the proteins that change significantly in both WCE and iPOND data (Group 2). E. Quantitative image-based cytometry (QIBC) analysis showing relative SETDB1 abundance in iPSCs and PSM-48-hour cells. Each dot represents the mean nuclear SETDB1 intensity per nucleus per well. The results are normalized to the level of expression in G1 phase iPSCs. Nuclei were gated into cell cycle phase according to their total DAPI intensity and mean EdU intensity. SETDB1 siRNA on all nuclei, not gated by cell cycle phase, is shown for reference. The results of three biological replicates are shown. F. As for E but shows ATF7IP relative mean nuclear intensity and the results of two independent biological replicates, are shown. See also Figure S3 and Table S3 and S4.

This analysis reveals that proteins recruited to nascent chromatin can be divided into 2 groups, stemming from distinct delivery pathways: one group where proteins are simply recruited to nascent chromatin upon pluripotency/differentiation signals (Group 1), and another group where proteins are expressed and subsequently recruited to nascent chromatin (Group 2). K-means clustering of these two groups independently revealed their various association patterns to replicated chromatin (Group 1: 321 proteins into Clusters I-V; Group 2: 182 proteins into Clusters VI and VII) (Figure 3C, S3B and Table S4). Each cluster is associated with distinct REACC terms (Figure S3C). STRING interaction network analysis of proteins enriched on nascent chromatin upon PSM induction (merged clusters II and VII) is shown in Figure 3D. Nodes outlined in black are proteins from Group 1 (already present in iPSCs and simply recruited to nascent chromatin upon differentiation), outlined in grey are proteins from Group 2 (proteins expressed and recruited to nascent chromatin upon differentiation). The largest network contains proteins associated with gastrulation including many TFs, and 25 of 30 are from Group 2. The second largest network contains proteins involved in heterochromatin assembly, and strikingly 11 of 16 proteins are from Group 1. Therefore, lineage specific TFs are expressed and recruited to newly replicated chromatin, while most heterochromatin assembly factors are already present in iPSCs and recruited to newly replicated chromatin upon a differentiation signal.

The next networks are Formation of SAHF, SUMOylation, and Fanconi anaemia pathway (Figure 3D). Further inspection of DNA repair proteins revealed that while FANC accumulate on nascent chromatin, level of SAMHD1, a dNTPase that handles stalled replication forks, decreases (Figure S3D and S3E). Importantly, measuring overall DNA damage level upon differentiation by immunofluorescence and by western blot using γH2AX shows an overall decrease in DNA damage level (Figures S2E, S3F), including in S phase cells (Figure S3G). Altogether this analysis shows that upon differentiation, although cells experience an overall reduced level of DNA damage, the replisome faces different challenges requiring distinct sets of repair proteins.

For proteins whose abundance is reduced on nascent chromatin upon differentiation, for group 2, the top REACC term detected is transcriptional regulation of pluripotent stem cells (Figure S3B, S3C, S3H). For group 1, the top REACC term is metabolism of RNA. Two of these RNA metabolism proteins WDR33, LSM3, may also have functions as putative DNA damage suppressors [54], while others are known as “sonication resistant” heterochromatin proteins: ERH, TRA2A, SLTM, SFAB, SAFB2, and CEBPZ [41]. Therefore, distinct heterochromatic proteins bind newly replicated chromatin in iPSCs and PSM cells, despite most remaining stably expressed.

The identification of a group of proteins already expressed in iPSCs and simply recruited to nascent chromatin upon differentiation, suggests that differentiation signals provoke or facilitate their recruitment to the replisome, acting as potential “first responders” to differentiation signals. To gain insight into how their recruitment may be regulated, we focused on two of these factors: SETDB1 and cofactor ATF7IP [55–58], and tested the hypothesis that differentiation signals promote SETDB1 import in the nucleus thereby facilitating its recruitment to newly replicated chromatin. By single-cell high-content microscopy we tracked changes in nuclear intensity of SETDB1 and ATF7IP in iPSCs and PSM cells. In iPSCs, SETDB1 nuclear level is reduced in S phase nuclei, to a similar level as siSETDB1 (Figure 3E). Upon PSM induction SETDB1 S-phase levels increased above the siRNA background level. Therefore, in response to PSM induction, the nuclear trafficking of SETDB1 is modified, enabling its retention in S phase cells. Interestingly, the nuclear intensity of ATF7IP also increased upon differentiation (Figure 3F) suggesting that ATF7IP may contribute to SETDB1 retention in the nucleus. Altogether these findings suggest that, in response to developmental cues, proteins can newly associate with nascent chromatin through mechanisms independent of transcriptional changes such as altering nuclear import upon S phase entry.

### Proteins selectively recruited to nascent chromatin in human iPSCs or in PSM cells are needed for differentiation

Finally, we examined the role of DNA replication in PSM induction and more specifically the role of distinct replisome interacting proteins identified in this study, SETDB1 and ERH. To examine the role of DNA replication in PSM differentiation we used two approaches (Figure 4A). Firstly, treatment with cell cycle inhibitors; Hydroxyurea (HU), which will prevent S phase progression, or palbociclib (PD), which will block cells in G1 phase (Figure 4B). In other differentiation systems, blocking G1 phase or S phase progression have impaired the silencing of pluripotency markers and/or the expression of lineage specific TFs [59–62]. Secondly, depleting CAF1a, thereby blocking DNA replication-coupled chromatin assembly (Figure 4C) that has been shown to impair the silencing of pluripotency markers and to provoke the binding of aberrant lineage TFs [12, 13, 17, 63, 64]. Following these treatments, we monitored level of pluripotency markers (SOX2, NANOG, and/or PRDM14) and the mesoderm lineage specific TFs (TBXT, TBX6, LEF1, CDX2) upon PSM induction using high content single cell microscopy and immunoblot.

**Figure 4:**
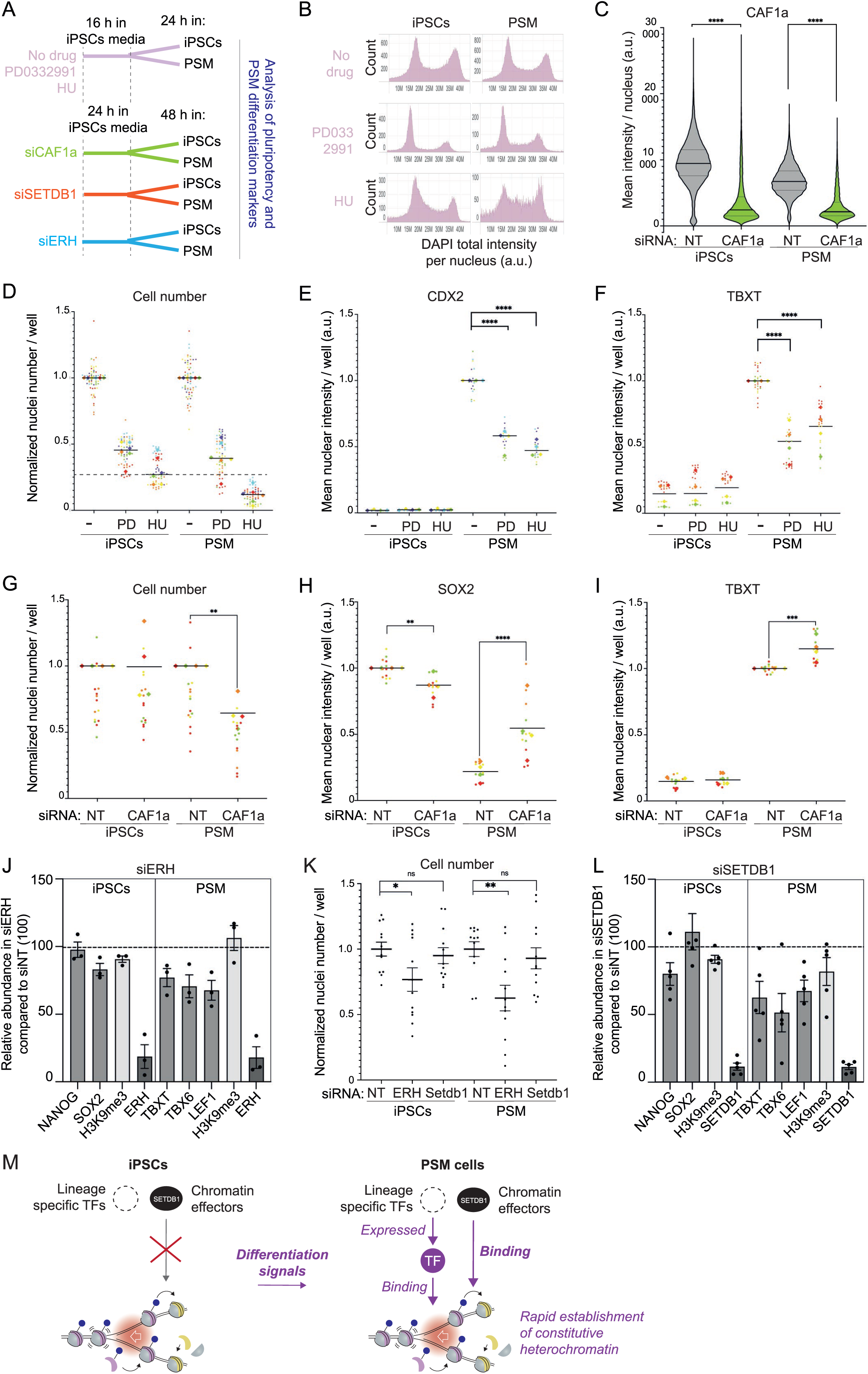
Knockdown of replisome associated proteins perturbs cell identity and differentiation. A. Schematic of the Hydroxyurea (HU) and palbociclib/PD0332991 (PD) treatments. Below, CAF1a, ERH or SETDB1 siRNA experiments including 24 hours siRNA treatment in iPSCs media followed by 48 hours +/- PSM differentiation. B. Cell cycle profile of iPSCs and PSM cells after PD and HU treatment. In QIBC experiments the total DAPI nuclear intensity was measured per nucleus after 40 hours PD/HU treatment. X axis shows the DAPI nuclear intensity and Y axis the number of nuclei with that intensity. C. Quantitative image-based cytometry (QIBC) analysis showing CAF1a (CHAF1A) levels in iPSCs and 48-hour PSM cells treated with non-target (NT) and CHAF1A (CAF1a) siRNA. D. Cell proliferation over 40 hours of treatment with PD and HU in iPSCs maintained as iPSCs or PSM differentiated for 24-hours. In QIBC experiments the number of nuclei per well were counted (n=6 biological replicates). E. QIBC analysis showing relative CDX2 nuclear intensity in iPS and PSM 24-hour cells that are untreated (-) or treated with PD or HU. The results of four biological replicates are shown. F. Same as D but relative TBXT mean nuclear intensity is shown. G. Proliferation upon CAF1a siRNA. In QIBC experiments the number of nuclei per well were counted. The results of four biological replicates are shown. H. QIBC analysis showing relative SOX2 nuclear intensity in iPSCs and PSM 48-hour cells with and without CAF1a siRNA. The results of four biological replicates are shown. I. Same as for G but TBXT/brachyury relative intensity is shown. J. Summary and quantification of three biological replicates of western blot experiments after ERH siRNA showing the protein abundance relative to the abundance in the non-target control siRNA cells. Each sample was also normalized to histone H3. Left; relative abundance in iPSCs, right; relative abundance in PSM cells. K. Nuclei number during ERH and SETDB1 siRNA. In QIBC experiments the number of nuclei per well were counted after 72 hours siRNA and plus/minus 48 hours of differentiation. The results of three biological replicates are shown. L. Same as for J but for SETDB1 siRNA. M. Model for replication coupled chromatin changes that drive differentiation. See also Figure S4.

PD and HU treatment blocked cell proliferation as expected (Figure 4B, 4D). Moreover, cells showed impaired expression of differentiation markers CDX2 and TBXT compared to the untreated control (Figure 4E, 4F). The expression of the pluripotency TF PRDM14 on the other hand was only slightly dysregulated (Figure S4A). This revealed that the G1/S transition and S phase progression facilitate the expression of PSM differentiation markers but maybe dispensable for pluripotency exit. Next, the level of the largest subunit of CAF1, CAF1a, was successfully reduced via siRNA (Figure 4C, 4G, S4B). Upon PSM induction, the pluripotency marker SOX2 was not properly downregulated upon siCAF1a depletion (Figure 4H), while the level of the mesoderm marker TBXT increased similarly in non-targeted siRNA (siNT) and siCAF1a (Figure 4I). Altogether these results reveal that cell cycle progression and DNA replication coupled chromatin assembly contribute to PSM induction and further confirm the important role of DNA replication in cell fate transitions.

Two proteins recruited or evicted from newly replicated chromatin upon differentiation, ERH and SETDB1, were next selected to test their contribution to PSM differentiation. Both proteins are stably present in iPSCs and PSM cells and have been linked to H3K9me3 which increases upon PSM induction (Figure 2H, S2D, S2E). Yet, ERH is enriched on iPSCs nascent chromatin (Figure 2D, S3H) while SETDB1is enriched on PSM-48-hour nascent chromatin (Figure 2D, 3D). Upon siRNA treatment, ERH expression was reduced three to tenfold across three independent biological replicates (Figure 4J, S4C). ERH knockdown caused negligible reduction in pluripotency marker expression and no aberrant increase in PSM markers in iPSCs (Figure 4J, S4D). However, upon PSM induction, expression of PSM markers TBXT, TBX6, and LEF1 are reduced, revealing that PSM induction was impaired. Of note, ERH affects proliferation of both iPSCs and PSM cells (Figure 4K). Therefore, we cannot exclude that the effect of ERH depletion on TBXT, TBX6 and LEF1, may be because of its impact on proliferation. SETDB1 expression was reduced five to tenfold by siRNA across three independent biological replicates and resulted in little change in iPSC pluripotency marker expression (Figure 4L, S4E). However, upon PSM induction, SETDB1 depletion reduced expression of PSM markers TBXT, TBX6, and LEF1, revealing that PSM induction was impaired (Figure 4L). Importantly, SETDB1 and ERH are known to regulate levels of H3K9me3 [41, 65–67], yet we observed no significant changes in H3K9me3 (Figure 4J, 4L), suggesting that ERH and SETDB1 may promote differentiation independently from H3K9me3 [56, 65]. Together these data revealed that CAF1 promotes the silencing of pluripotency markers during PSM induction. The expression of mesoderm specific TFs on the other hand relies on cell cycle progression through S-phase and the presence of ERH, and SETDB1. Moreover, this suggests that proteins rapidly recruited to or evicted from nascent chromatin in response to differentiation signals contribute to cell fate transition.

## Discussion

Here we uncover extensive remodelling of replicated chromatin upon differentiation from iPSCs to PSM cells. Specifically, we observed dynamic alterations in TF composition, epigenetic modifiers, and histone chaperones on newly replicated chromatin, (Figure 4M), with over one hundred proteins newly recruited to PSM nascent chromatin (Figure 1G). Notably, the dynamic recruitment of many of these proteins is undetectable by whole cell based proteome analysis (Figure 3C) and studies using RNA-seq [27, 32]. Thus, our findings identify a novel subset of proteins, revealing an understudied aspect of differentiation-associated chromatin dynamics. Previous studies have indicated early changes, such as dephosphorylation, precede transcriptional onset suggesting transcription-independent mechanisms can regulate the earliest stages of differentiation [68]. As most proteins recruited to nascent chromatin upon differentiation were already present in pluripotent cells, we propose they have a role as first responders to differentiation signals, and this raises new important questions about how differentiation cues instruct the selective recruitment of these pre-existing chromatin factors.

Multiple genome-wide studies have documented changes in histone modifications accompanying differentiation, notably resolution of bivalent domains, and increase in specific histone marks such as H3K9 methylation [42, 66, 69–73]. For example, genome-wide increases in H3K9me3 occur during germ-layer development and gastrulation and genes enriched for H3K9me3 typically exhibit reduced TF accessibility, safeguarding cellular identity by suppressing inappropriate lineage-specific transcription [41, 42, 66, 72, 73]. However, comprehensive characterization of replication-coupled histone modifications during differentiation has never been undertaken. Our study, for the first time, detects histone modifications dynamics on newly replicated chromatin upon differentiation. Strikingly there is an increase in the heterochromatin-associated H3K9me2 and H3K9me3 on nascent chromatin during differentiation (Figure 2H). Elevated H3K9 methylation upon differentiation may be initiated directly on nascent chromatin or reinforced rapidly post-replication, thereby limiting alternative lineage activation and/or repressing retrotransposons during lineage commitment. Consistent with this on PSM nascent chromatin we see increased association of DAXX; recently shown to deliver H3K9me3 to newly replicated chromatin [74]. Moreover, from 24 hours post-PSM induction, we detect increased association of H3K9me3 methyltransferase SETDB1, and its cofactor ATF7IP with newly replicated chromatin, as well as the demethylases KDM4B (Figure 2D). These factors may have complementary roles: SETDB1 in repressing inappropriate lineage programs and KDM4B in promoting the activation of lineage-specific genes [42, 66]. ERH and SAFB, both associated with H3K9me3 and known repressors of satellite repeats [41, 75], were preferentially enriched on nascent chromatin in undifferentiated iPSCs (Figure 2D, S3H). This suggests that repeat repression mechanisms mediated by ERH operate immediately after DNA replication in pluripotent states but not in differentiated state. Given the central role of ERH and SETDB1 in H3K9 methylation, it was unexpected that knockdown of either protein did not result in a global loss of H3K9me3 (Figure 4J, 4L). This may reflect localized reductions in H3K9me3 restricted to nascent chromatin, which may not be detectable in bulk analyses, or alternatively, that ERH and SETDB1 initially influence differentiation independently of their role in H3K9 methylation. Evidence from other systems supports the latter possibility [56, 65].

Finally, CAF1 is essential for maturation of nascent chromatin and for preservation of lineage fidelity during differentiation [11, 64, 76]. Our results reinforce that indeed progression through S-phase and CAF1-dependent chromatin assembly are critical for successful differentiation of iPSCs into PSM cells (Figure 4). Both TFs and CAF1 can limit chromatin accessibility immediately following DNA replication but newly replicated chromatin is still hyperaccessible, providing a temporal window for TF binding [11]. However, accessibility alone does not fully explain accurate TF engagement [77]. Other factors such as post-translational modifications, motif strength, and interaction partners may also be important. Further experiments (e.g., CHOR-seq or SCAR-seq) will be necessary to define how TFs and other factors precisely engage their genomic targets post-replication.

Collectively, our data suggest a model where replication-coupled chromatin assembly plays a central role in defining cellular identity and lineage commitment. Since most proteins newly recruited upon differentiation were already expressed in pluripotent cells, our works identifies new previously overlooked targets that may be important for differentiation onset. Future work will uncover if their recruitment specifically to newly replicated chromatin is essential for differentiation. It is likely that replication-associated chromatin hyperaccessibility might facilitate their rapid mobilization in response to developmental signals but precisely how remains to be elucidated.

### Limitations

Our proteomics analyses do not allow resolution at specific genomic loci and were performed at the population level, potentially obscuring cell-to-cell variability. For instance, at the 24-hour differentiation time point, cellular heterogeneity may mask specific chromatin changes (Figure 1B, S1D, and S1E). Single-cell and FUCCI-based approaches could address these limitations by clarifying factor-specific cell cycle recruitment dynamics at this differentiation stage. Additionally, the asynchronous nature of the analyzed cell population complicates interpretation of chromatin changes occurring later on mature chromatin.

While we identify factors; ERH, SETDB1, and CAF1, enriched on nascent chromatin that are essential for differentiation, subsequent functional assays employing siRNA-mediated knockdowns do not allow knockdown only at nascent chromatin, complicating mechanistic interpretations. Currently, technical limitations prevent protein manipulation only at nascent chromatin, necessitating future technology development. Furthermore, our validation of ERH, SETDB1, and CAF1a focused on a limited set of pluripotency and differentiation markers, potentially underestimating their overall roles. Comprehensive, large-scale analyses, such as RNA-seq or MS, following knockdown, could further elucidate the broader impacts of these factors on differentiation.

## Resource Availability

### Lead Contact

Requests for further information and resources should be directed to Constance Alabert

### Materials availability

All reagents are available upon request

### Data and code availability

The mass spectrometry proteomics data have been deposited to the ProteomeXchange Consortium via the PRIDE partner repository and are publicly available as of the date of publication. Accession numbers are listed in the key resources table.

This paper does not report original code, but any code or additional information required to reanalyse the data reported in this paper is available from the lead contact upon request.

## Acknowledgments

This work was predominantly funded by the European Research Council ERC-Stg-IDRE. The authors thank Margarita Kalamara from the Human Pluripotent Stem Cell Facility and the Imaging facility, both University of Dundee, for their contributions to this work. We acknowledge experimental support from Pei Rong Toh and Nadia Lefter. We would like to thank Kate Storey and Hedda Meijer for advice and comments on the manuscript. Research in the Lamond lab was funded by BBSRC grant (Ref: BB/V010948/1) and UKRI grant (Ref: EP/Y010655/1). Research in the Alabert lab was funded by CRUK-CDF (C57404/A21782) and the European Research Council ERC-Stg-IDRE.

## Author contributions

Conceptualization, J.E.W., and C.A.; methodology, J.E.W., C.A, S.B., H.J., S.D.S, L.D, M.V.A, and A.L.; validation, J.E.W.; formal analysis, J.E.W., H.J., S.D.S, and M.V.A; investigation, J.E.W., S.B., H.J., and S.D.S.; writing – original draft, J.E.W. and C.A.; writing – review & editing, J.E.W., and C.A. ; funding acquisition, C.A. and A.L.

## Declaration of interests

The authors declare no competing interests.

## METHODS

### KEY RESOURCES TABLE

| REAGENT or RESOURCE | SOURCE | IDENTIFIER |
| --- | --- | --- |
| **Antibodies** |  |  |
| NANOG | Cell signalling technology | Cat#CST4903 |
| SOX2 | Cell signalling technology | Cat#CST3579 |
| PRDM14 (E1D5S) | Cell signalling technology | Cat#CST83527 |
| TBXT | R and D | Cat#AF2085 |
| CDX2 (D11D10) | Cell signalling technology | Cat#CST12306 |
| LEF1 | Cell signalling technology | Cat#CST2230 |
| TBX6 | R and D | Cat#AF4744 |
| ERH [EPR10830(B)] | Abcam | Cat#ab166620 |
| SETDB1 (ESET) C1C12 | Cell signalling technology | Cat#CST2196 |
| CHAF1A D77D5 mAb | Cell signalling technology | Cat#CST5480 |
| CAF1 p150 | Bethyl | Cat#A301-481A-M |
| PCNA (Mouse Monoclonal, Clone PC10) | Abcam | Cat#ab29 |
| Ubiquityl-Histone H2B (Lys120) (D11) | Cell signalling technology | Cat#CST5546 |
| Pan-H3 | Abcam | Cat#ab10799 |
| Ubiquityl-Histone H2A (Lys119) (D27C4) | Cell signalling technology | Cat#CST8240 |
| Phospho-Histone H2A.X (Ser139) | Cell signalling technology | Cat#CST2577 |
| H3K27me3 | Cell signalling technology | Cat#CST9733 |
| H3K9me3 | Abcam | Cat#AB176916 |
| **Chemicals, peptides, and recombinant proteins** |  |  |
| Palbociclib PD-0332991 | Sigma-Aldrich | Cat#PZ0383 |
| Hydroxyurea (HU) | Sigma-Aldrich | Cat#H8627 |
| 5-ethynyl-2'-deoxyuridine (EdU) | Thermo Fisher Scientific | Cat# E10187 |
| Thymidine | Sigma-Aldrich | Cat#T1895 |
| Biotin-azide | Thermo Fisher Scientific | Cat# B10184 |
| Alexa Fluor™ 647 Azide, Triethylammonium Salt | Thermo Fisher Scientific | Cat#A10277 |
| Streptavidin-coated magnetic beads | Thermo Fisher Scientific | Cat# 65001 |
| Geltrex basement membrane matrix | Thermo Fisher Scientific | Cat#A1413302 |
| Recombinant human FGF-2 | PeproTech | Cat# 100-18B |
| Recombinant human Noggin | PeproTech | Cat# 120-10C |
| TrypLE Select (1X) | Thermo Fisher Scientific | Cat#12563011 |
| ROCK inhibitor (Y-27632) | TOCRIS | Cat#1254 |
| CHIR99021 | TOCRIS | Cat#4953 |
| LDN193189 | Sigma-Aldrich | Cat#SML0559 |
| **Critical commercial assays** |  |  |
| TMT10plex™ Isobaric Mass Tag Labeling Kit | Thermo Fisher Scientific | Cat# 90110 |
| Click-iT® EdU Imaging Kit | Thermo Fisher Scientific | Cat# B10184 |
| TeSR1 medium | In-house; also available from STEMCELL Technologies | N/A |
| **Deposited data** |  |  |
| iPOND MS of iPSCs, 24 and 48 hours PSM (no EdU, N, and M) | This study | PRIDE: PXD059836 |
| Whole cell extracts MS of iPSCs, 24 and 48 hours PSM differentiation | This study | PRIDE: PXD059790 |
| Histone PTMs | This study | PRIDE: PXD061922 |
| **Experimental models: Cell lines** |  |  |
| ChiPSC4 (Cellartis human induced Pluripotent Stem Cell line 4) | Cellartis/Takara Bio Europe or Human Pluripotent Stem Cell Facility, University of Dundee | CVCL_RM97 |
| **Oligonucleotides** |  |  |
| On-target plus Non-targeting Control Pool | Horizon discovery | D-001810-10 |
| On-target plus CHAF1A siRNA | Horizon discovery | L-019938 |
| ERH silencer select siRNA | Thermo Fisher Scientific | s4814 |
| ERH silencer select siRNA | Thermo Fisher Scientific | s4815 |
| SETDB1 silencer select siRNA | Thermo Fisher Scientific | s19110 |
| SETDB1 silencer select siRNA | Thermo Fisher Scientific | s19111 |
| **Software and algorithms** |  |  |
| MaxQuant (v 1.6.10.43) | [78] | https://maxquant.net/maxquant/ |
| Perseus (1.6.7.0) | [79] | https://maxquant.net/perseus/ |
| RStudio (R 4.3.1) |  | https://www.r-project.org |
| PRISM | GraphPad | https://www.graphpad.com/scientific-software/prism/ |
| g:Profiler | [80] | https://biit.cs.ut.ee/gprofiler/gost |
| STRING | [81] | https://string-db.org |
| CYTOSCAPE (3.10.3) | [82] | https://cytoscape.org |
| **Other** |  |  |
| ScanR High Content Screening Microscopy | Olympus | N/A |
| Orbitrap Fusion™ Tribrid™ Mass Spectrometer | Thermo Fisher Scientific | N/A |
| Licor FC |  |  |

### EXPERIMENTAL MODEL AND STUDY PARTICIPANT DETAILS

Cellartis human induced Pluripotent Stem Cell line 4 (ChiPSC4) cells were maintained in TESR medium [83] containing FGF2 (30 ng/ml) and noggin (10 ng/ml) on growth factor reduced geltrex basement membrane extract (10 μg/cm^2^) coated dishes at 37 °C in a humidified atmosphere of 5% CO2 in air. Media was changed daily when cells not passaged. Cells were passaged twice a week as single cells using TrypLE select and replated in TESR medium that was further supplemented with the Rho kinase inhibitor Y27632 (10 μM). Twenty-four hours after replating, the media is replaced with media without Y27632 (mTESR with FGF and noggin). ChiPSC4 cells were differentiated along the pre-somitic mesoderm lineage by removing TESR medium (with FGF, noggin and Y27632), washing in DMEM/F12 (Thermo Fisher 21331-046), and replacing with DMEM/F12 medium supplemented with Glutamine, Non-essential amino acids (NEAA; Thermo Fisher 11140035), 1% insulin–transferrin–selenium (ITS; Thermo Fisher 41400045), 3 μM CHIR99021, and 0.5 μM LDN193189 for 24, 48 or 72 hours [26, 27]. Media was replenished daily.

### METHOD DETAILS

#### Drug treatments

Hydroxyurea (HU: 300 μM), DNA synthesis inhibitor, or PD0332991 (PD: 5 μM), selective inhibitor of cyclin kinases CDK4 and CDK6, were added to iPSCs in TESR medium (with FGF, noggin) for 16 hours and then cells were either replenished with fresh media containing HU/PD or were washed in DMEM/F12 and PSM media (DMEM/F12 +Glutamine, 1% NEAA, 1% ITS, 3 μM CHIR99021, and 0.5 μM LDN193189) containing HU/PD added for a further 24 hours.

#### DNA labelling

For iPOND and Imaging-based cell cycle analysis cells were labelled with medium containing EdU at a final concentration of 40 μM for 20 min. For imaging, cells were harvested immediately. For iPOND, following labelling, nascent samples were immediately taken and processed and mature samples were washed twice in 1× DMEM/F12 and further incubated in fresh media containing 40 μM thymidine for either three or eleven hours before collection.

#### Isolation of proteins in nascent DNA (iPOND)

iPOND was performed essentially as described previously [23, 25]. Briefly, per time point, 1 × 10^8^ asynchronous cells were EdU-labelled as described above and then cells were crosslinked with 1% formaldehyde for 15 min at RT followed by 5-min incubation with 0.125 M glycine. Cells were then scraped from plates on ice, permeabilized in 0.25% Triton X-100 in PBS for 30 min at RT and washed with 0.5% BSA in PBS. To conjugate biotin to EdU-labelled DNA, click chemistry reactions were performed for 1 hour in the dark at RT (10 μM Biotin-azide, 10 mM Sodium ascorbate, 2 mM CuSO4 in 1×PBS). Cells were lysed (1% SDS in 50 mM Tris, pH 8.0, protease inhibitor), sonicated (Bioruptor, Diagenode), the lysate diluted 1:1 (v/v) with cold PBS containing protease inhibitors. At this point, aliquots of the lysate were collected to be analysed as whole cell extract (WCE). The majority of the lysate was incubated overnight at 4 ^0^C with streptavidin beads to capture the biotin-conjugated DNA-protein complexes. Captured complexes were washed using lysis buffer and 1 M NaCl. The proteins were eluted under reducing conditions by boiling in 2× LSB sample buffer for 30 min.

#### Tandem mass tag-based MS quantitative proteomics analysis (TMT MS) of iPOND and WCE

Samples were prepared using SP3 protocol for TMT labelling as described in [84]. In brief, both iPOND and Whole Cell Extract (WCE) samples were denatured at 95 ℃ for 10 min, alkylated with 400 mM IAA (final concentration) in dark at RT for 30 min. Protein concentration was measured using EZQ™ Protein Quantitation Kit. Then, protein samples were mixed and cleaned with SP3 beads (1: 10, protein:beads), and digested with lysC/trypsin mixture (1: 50, enzyme:protein). 100 μL 100 mM TEAB buffer (pH 8.0) was used to elute the peptides from SP3 beads. Peptides from each sample were labelled with TMT11plex Isobaric Mass Tag Labelling Kit (Thermo Fisher Scientific). For WCE samples, 100 ug peptides from each sample were labelled. In addition, a pool of peptides from all WCE samples were created, labelled with one TMT channel and added as the reference channel for batch normalisation. TMT labelled peptide samples were further fractionated using offline high-pH reverse-phase (RP) chromatography, as previously described [85]. The 24 fractions were subsequently dried, and the peptides re-dissolved in 5% formic acid and analysed by nanoLC–MS/MS system Orbitrap Fusion Tribrid Mass Spectrometer (Thermo Fisher Scientific), equipped with a Dionex ultra-high-pressure liquid-chromatography system (RSLCnano). RPLC was performed using a Dionex RSLCnano HPLC (Thermo Fisher Scientific). Peptides were injected onto a 75 μm × 2 cm PepMap-C18 pre-column and resolved on a 75 μm × 50 cm RP-C18 EASY-Spray temperature-controlled integrated column-emitter (Thermo Fisher Scientific), using a 120-min multistep gradient from 10% B to 44% B with a constant flow rate of 300 nl/min. The mobile phases were H2O incorporating 0.1% FA (solvent A) and 80% ACN incorporating 0.1% FA (solvent B). The data were acquired under the control of Xcalibur software in a data-dependent acquisition mode (DDA) using top speed and 3 second duration per cycle. The survey scan was acquired in the orbitrap covering the m/z range from 375 to 1,600 Da with a mass resolution of 120,000 and a stand automatic gain control (AGC) target. The most intense ions were selected for fragmentation using CID in the ion trap with 35% CID collision energy and an isolation window of 0.7 Da. The AGC target was set to standard with automatic maximum injection time and a dynamic exclusion of 60 s. The MS2 scan was acquired in the ion trap in Turbo mode. SPS-MS3 mode was applied for more accurate TMT quantifications. 10 fragment ions were co-isolated using synchronous precursor selection with a window of 3 Da and further fragmented using HCD collision energy of 65%. The fragments were then analyzed in the orbitrap with a resolution of 50,000. The AGC target was set to standard, and the maximum injection time was set to 105 ms.

The TMT-MS data were analysed using MaxQuant (v 1.6.10.43, [78]) and searched against Homo sapiens database from Uniport (Swiss-Port only, downloaded in October 2021). The FDR threshold was set to 1% for each of the respective Peptide Spectrum Match (PSM) and Protein levels. The data was searched with the following parameters: quantification type was set to Reporter ion MS3 with TMT 11plex, stable modification of carbamidomethyl (C), variable modifications, oxidation (M) and acetylation (protein N terminus), with maximum of 2 missed tryptic cleavages threshold. The detailed parameter file was uploaded together with the raw MS data.

#### Sample preparation for histone modification analysis by MS

The iPOND was performed as described before with minor changes. There was an additional mature chromatin timepoint of 11 hours. Captured complexes on streptavidin beads were washed with lysis buffer and 1 M NaCl, followed by two washes with 50 mM Tris, pH 8 and then dried bead pellets submitted for further analysis. For whole cell extracts, iPSCs and cells differentiated along PSM pathway for 24 or 48 hours, were harvested from dish using TrypLE select, washed in media, washed in PBS, cells counted, and then cell pellets flash frozen. Approximately 1 million cells were used and acid extracted histones were processed according to a SP3 protocol as described previously [86]. However, proprietary steps developed by MOLEQLAR Analytics GmbH have been added in order to adjust the protocol for histone specific aspects. For iPOND samples already provided as dried beads, the protocol started from the protein propionylation. Upon 3 hours digestion at 37 °C and 1,000 rpm in a table-top thermomixer, samples were acidified by adding 5 µl of 5% TFA and quickly vortexed. Beads were immobilized on a magnetic rack, and peptides were recovered by transferring the supernatant to new tubes. Samples were dried down using a vacuum concentrator and reconstituted by adding 20 µl 0.1% FA to reach a peptide concentration of approximately 0.2 µg/µl. MS injection-ready samples were stored at -20 °C.

#### LC-MS analysis of histone modifications

Approximately 200 ng of peptides from each sample were separated on a C18 column (Aurora Elite TS, 15 cm x 75 μm ID, 1.7 μm, IonOpticks) with a gradient from 5% B to 25% B (solvent A 0.1% FA in water, solvent B 100% ACN, 0.1% FA) over 35 min at a flow rate of 300 nl/min (Vanquish Neo UHPLC-Systems, Thermo - Fisher, San Jose, CA) and directly sprayed into a Exploris 240 mass spectrometer (Thermo-Fisher Scientific). The mass spectrometer was operated in full-scan mode to identify and quantify specific fragment ions of N-terminal peptides of human histone 3.1 and histone 4 proteins. Survey full scan MS spectra (from m/z 250–1600) were acquired with resolution 60,000 at m/z 400 (AGC target of 3×10^6^). Typical mass spectrometric conditions were: spray voltage, 1.9 kV; no sheath and auxiliary gas flow; heated capillary temperature, 300 °C.

#### Quantification of histone modifications

Data analysis was performed with Skyline (version 23.1.0.455) [87] by using doubly and triply charged peptide masses for extracted ion chromatograms (XICs). Peaks were selected manually. Heavy arginine-labelled spiketides (13C6; 15N4) were used to confirm the correct retention times and for signal normalization purposes, because all heavy standards were incorporated across all samples at the same concentration. Integrated peak values (Total Area MS1) were used for further calculations. Endogenous PTM signals were normalized according to the variation of the signals of the spiked-in heavy standards and using median normalization. The percentage of each modification within the same peptide is derived from the ratio of this structural modified peptide to the sum of all isotopically similar peptides. Therefore, the Total Area MS1 value was used to calculate the relative abundance of an observed modified peptide as percentage of the overall peptide. The unmodified peptide of histone 3.1 (aa 41–49) was used as indicator for total histone 3.1. Coeluting isobaric modifications were quantified using three unique MS2 fragment ions. Averaged integrals of these ions were used to calculate their respective contribution to the isobaric MS1 peak (e.g., H3K36me3 and H3K27me2K36me1).

#### SDS PAGE and Immunoblotting

Proteins were resolved in 4–12% Bis-Tris NuPAGE gels (Life Technologies, NP0301) in 1× MES buffer and then transferred onto 0.2-μm pore size nitrocellulose membranes (GE Healthcare, 10600001) using Trans-Blot SD Semi-Dry Transfer Cell (Bio-Rad). Membranes were blocked in 5% dried milk in TBST buffer (20 mM Tris base, 137 mM Sodium chloride pH 7.6, 0.1% Tween 20) for 1 hour at room temperature. Primary antibodies were diluted 1:1000 in 5% BSA in TBST and then incubated with membranes overnight at 4 °C. The membranes were then washed three times for 5 minutes with TBST, before incubation with secondary antibody (1:10000) with 5% dried milk in TBST, for 1 hour at room temperature. The membranes were washed again with TBST three times for 5 min, once with TBS and proteins detected on the Odyssey FC imaging system (LICOR).

#### Transfections with siRNA

siRNAs were introduced into iPSCs in TESR medium containing FGF2 (30 ng/ml), noggin (10 ng/ml), and Y27632 (10 μM) using Lipofectamine RNAiMAX (Invitrogen) according to manufacturer’s recommendations. The siRNAs used are shown in the key resources table and were used at the following final concentration; Non-Target Control (10 nM or 20 nM); CHAF1A (20 nM); ERH (10 nM); and SETDB1 (10 nM). Twenty-four hours after transfection, the media is either replaced with media without Y27632 or cells are washed in DMEM/F12 and PSM media (DMEM/F12 +Glutamine, 1% NEAA, 1% ITS, 3 μM CHIR99021, and 0.5 μM LDN193189) is added.

#### Immunofluorescence

For quantitative image-based cytometry (QIBC), cells were seeded from 10,000 to 20,000 cells/cm^2^ onto clear bottom black 96-well plates (Greiner), pre-coated with geltrex. At the end of each experiment, cells were fixed in 4% formaldehyde in PBS for 10 min, permeabilized with 0.25% triton in PBS for 5 minutes at 4 ^0^C. Cells were then washed in PBS, blocked with BSA and then either EdU detection and/or antibody detection was performed. For EdU detection, Alexa 647 was covalently linked to EdU using the Click-iT EdU Imaging Kit. For antibody detection, antibodies were incubated at 1:200 dilution except for: ERH (1:100); Phospho-Histone H2A.X Ser139 (1:800); anti-rabbit IgG AF488 (1:1000, 1910751, Invitrogen), and anti-Alexa Fluor 647 anti-goat IgG (H + L) (1:1000, A32849, Invitrogen). For DNA staining, the cells were incubated with DAPI (2.5 ug/ml Thermo Fisher Scientific, 62248) for 30 min at RT. The cells were then washed twice in PBS-Tween, one time with PBS, and left in PBS at 4 ^0^C until imaging. Plates were imaged on Olympus Scan-R imaging system using a 20x objective. 25 fields per well were imaged. Single-cell fluorophore intensities were extracted using Scan-R analysis software.

### QUANTIFICATION AND STATISTICAL ANALYSIS

#### Mass spectrometry data analysis for iPOND and WCE samples

MaxQuant software was used for the identification of proteins. MaxQuant’s output files, proteinGroups.txt, underwent analysis using Perseus and Rstudio. First, both iPOND and WCE datasets were filtered, removing proteins identified as reverse sequences, potential contaminants, and only identified by site.

For iPOND samples, to control for technical variation between the different batches, a correction factor, cf, was applied to every protein, in every batch, to adjust the intensity values. This cf was calculated per protein using the reference channel for batch normalisation. Reference channel was generated from a pool of peptides from all WCE samples, labelled with one TMT channel. Subsequently, median normalization was performed by dividing each column by its median. All replicates were grouped into their biological groups and filtered such that a protein must appear at least 3/3 times in any one group to be retained (4279 proteins). Dataset was further filtered for proteins with at least 2 unique peptides (4171 proteins) and then proteins more abundant in the negative control (no EdU labelling, +ClickIT: log_2_ >0.58 and p adjust <0.05) compared to the EdU labelled samples were removed from analysis since considered as background/non-specific proteins. (2956 proteins retained). To mitigate batch effects, we employed HarmonizR, using COMBAT method and any missing values were also replaced using HarmonizR [88]. Subsequent PCA verified substantial attenuation of batch effects.

For WCE samples, median normalization was performed by dividing each column by its median. All replicates were grouped into their biological groups and filtered such that a protein must appear at least 3/3 times in any one group to be retained. Dataset was further filtered for proteins with at least 2 unique peptides. Missing values were replaced using imputation in Perseus.

For determining differential enrichment, in both iPOND and WCE datasets, we used LIMMA package [89]. Volcano plots were generated in Rstudio. Significant changes were determined to be proteins exhibiting changes in pairwise contrasts determined by a false discovery rate of less than 0.05 and a log_2_ fold change of at least 0.9. Clustering (K-means) was performed on the 503 significant iPOND proteins. Individual Heatmaps were generated in GraphPrism using the row mean normalized log_2_ intensity values to visualize intensity variations across different conditions. Reactome (REACC) analyses were performed using gProfiler [80]. GraphPrism was used to visualise the results. STRING protein interaction networks were imported into Cytoscape [81, 82]. Only high confidence interactions were considered and MCL clustering performed with a granularity parameter of 3.

#### Mass spectrometry data analysis for histone modifications

The resulting peptide lists were exported to Microsoft Excel and further processed for a detailed analysis. Out of 35 detected modifications, 26 were analysed further based on data availability for of all modified state, and an abundance >1%. To calculate the relative abundance of post-translational modifications (PTMs), the sum of all different modified forms of a histone peptide was considered as 100%. Each particular modified peptide was calculated by dividing its intensity by the sum of all the modified form of the peptide. For peptides that contained more than one modification (e.g. H3K27me1 on H3 aa 27-40 peptide than also contain H3K36 methylations), the sum of all the possible combinations were considered (e.g. H3K27me1 = H3K27me1K36me0 + H3K27me1K36me1 + H3K27me1K36me2 + H3K27me1K36me3 + H3K27me1K36ac).

#### Western blot quantification

Image processing and band intensity quantification were performed using Odyssey FC imaging system (LICOR) and Image Studio software (LICOR). Reported band intensities were first normalized to pan-H3 loading control for each sample and then the relative protein abundance for treated samples shown relative to the control NT siRNA sample.

#### High throughput microscopy data analysis or Quantitative Image based Cytometry (QIBC)

Microscopy images were analysed firstly with ScanR analysis software and single-cell fluorophore intensities were extracted using Scan-R analysis software. All data were subsequently visualised and statistically analysed in Tableau Desktop 2023.2 and GraphPrism. Graphs were generated using Tableau and GraphPad Prism. Cell cycle phases were gated based on DAPI and EdU intensities.

#### Statistical analysis

For Figures 2, S2, 3, S3, all bar charts show averages and standard deviation calculated in GraphPrism and t tests applied were unpaired and two-sided using GraphPrism. For Figures 4 and S4 the superplots are colour coded to indicate the biological replicates and the P-values represent a paired two-tailed T-test. P values are indicated by asterisks (P < 0.0001 [****], P < 0.001 [***], P < 0.01 [**], and P < 0.05 [*]), and n.s. indicates nonsignificant.

---

**Figure S1.**
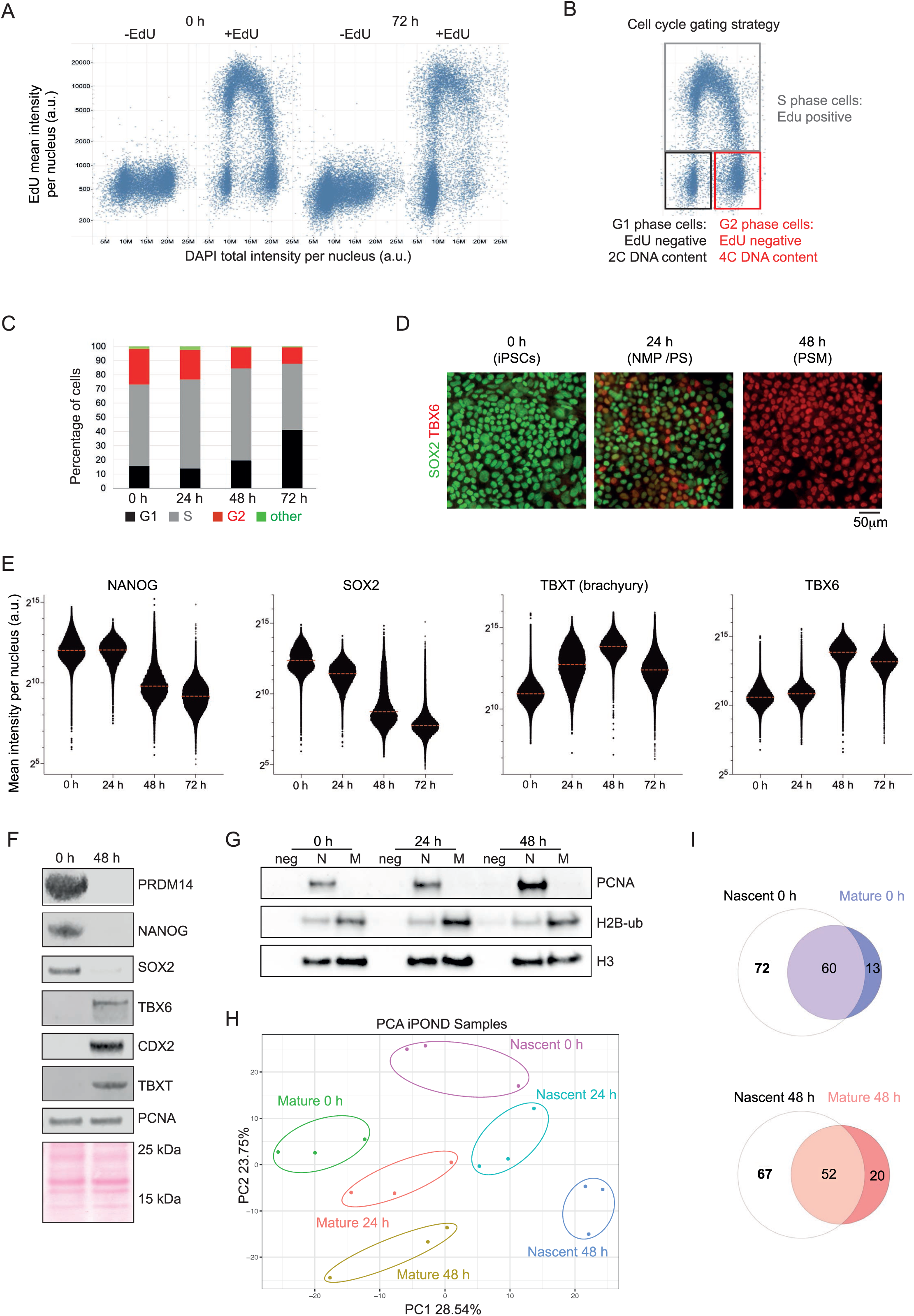
The presomitic mesoderm pathway is a suitable system to study the contribution of DNA replication associated mechanisms towards cell fate changes. A. Representative QIBC EdU/DAPI scatter plots to show the cell cycle distribution between iPSCs and cells differentiated for 72 hours along the PSM pathway. B. Cell cycle gating strategy. Cells were gated from QIBC EdU/DAPI scatter plots. EdU positive cells were gated as S phase cells (grey) and cells with low EdU and low DAPI gated as G1 phase (black) and low EdU, high DAPI gated as G2 phase (red). Cells not falling into these gates were termed other (green). C. Bar chart showing the percentage of cells in each cell cycle phase of iPSCs and PSM cells differentiated for 24, 48, or 72 hours. D. Representative immunofluorescence images to reveal levels of a pluripotency marker (SOX2) in green and a pre-somitic mesoderm marker (TBX6) in red at 0, 24, and 48 hours of PSM differentiation. E. Quantitative image-based cytometry (QIBC) analysis showing the nuclear mean intensity expression level of four proteins (NANOG, SOX2, TBXT, and TBX6) at 0, 24, 48, and 72 hours of differentiation along the PSM pathway. F. Representative western blot of various marker proteins contrasting iPSCs and 48-hour PSM-differentiated cells. G. Representative western blot of an iPOND experiment in iPSCs (0 h) and cells differentiated in PSM media for 24 and 48 hours. Cells were untreated (neg) or treated with EdU for 20 minutes and harvested immediately (nascent, N) or after 3 hours (mature, M). H. Principal component analysis (PCA) of three iPOND TMT MS biological replicates based on the proteins identified. PC1 versus PC2 is shown. List of significantly changing proteins is available in Table S1. I. Venn diagrams overlapping the proteins, shown in Figure 1G and 1H, who significantly change in nascent and mature chromatin between iPSCs and PSM cells. Left: Contrasting nascent and mature enriched proteins from iPSCs. Right: Contrasting nascent and mature enriched proteins from PSM 48-hour cells.

**Figure S2.**
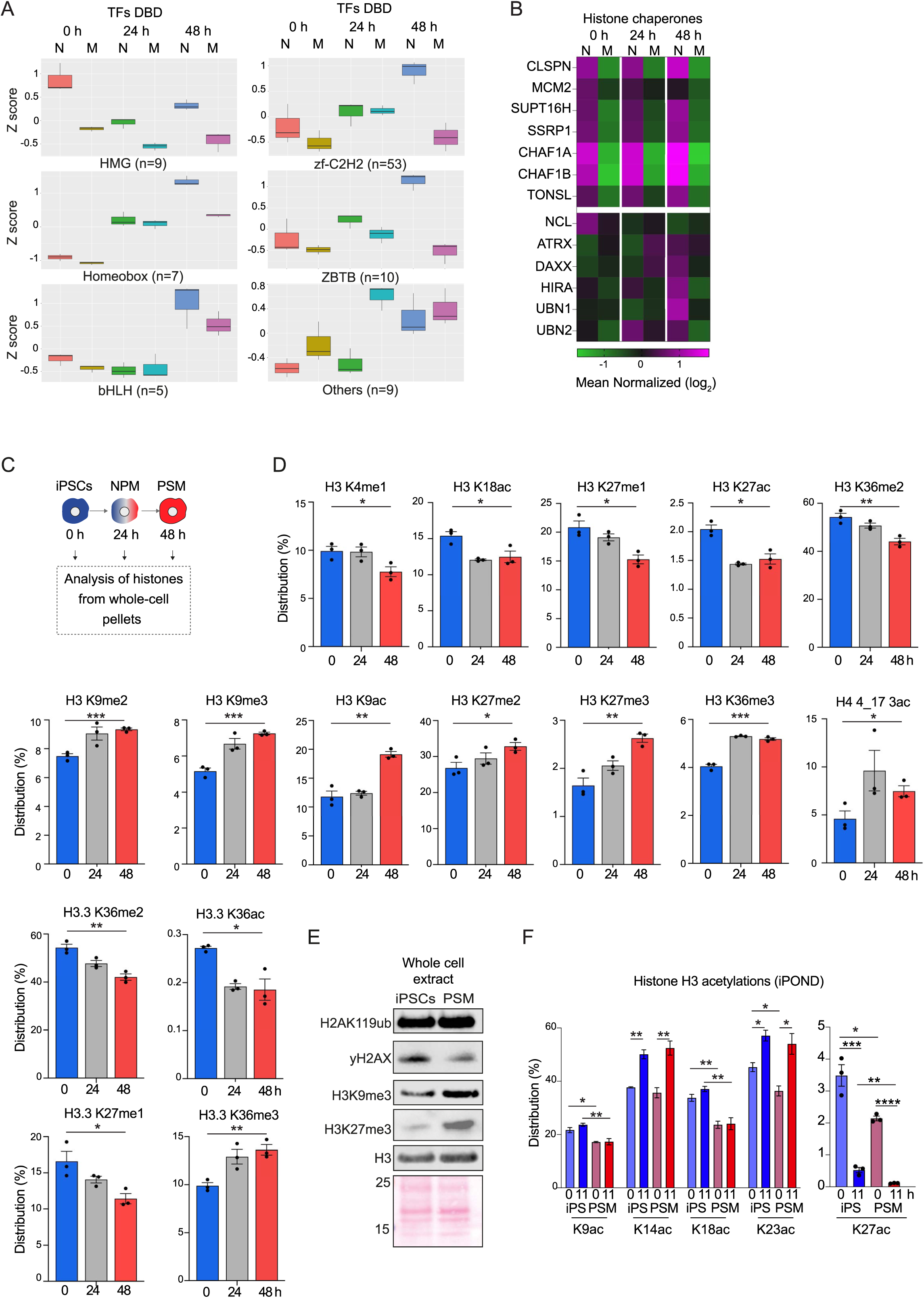
TFs and histones show different recruitment dynamics to replicated chromatin in iPSCs versus PSM cells. A. Each TF was categorized according to its main DNA binding domain (DBD) based on the TFDB4.0 database. For DBD groups that contain at least 5 proteins, their group metagene expression is shown. B. Heatmap showing differential binding of histone chaperone proteins to nascent and mature chromatin iPOND samples during PSM differentiation. Scale is mean normalized log_2_ (reporter intensities). C. Schematic for the presomitic mesoderm (PSM) differentiation pathway and the isolation of histones from whole cell pellets by acid extraction (see method). D. Dynamics of Histone H3/H4 modifications in WCE from iPSCs, PSM 24-hour, and PSM 48-hour cells. Only modifications that significantly change with differentiation are shown. Each modification is shown as a percentage of the relevant peptide detected. Percentages for each modification are available in Table S2. E. Representative western blot of various histone modifications contrasting total whole cell extract of iPSCs and 48-hour PSM-differentiated cells. F. Dynamics of Histone H3 K9, K14, K18, K23, and K27 acetylation on replicated chromatin (iPOND) in iPSCs and PSM 48-hour cells. H3 acetylation in nascent (0 h) and mature chromatin (+11 h) is shown as a percentage of the relevant peptide detected.

**Figure S3.**
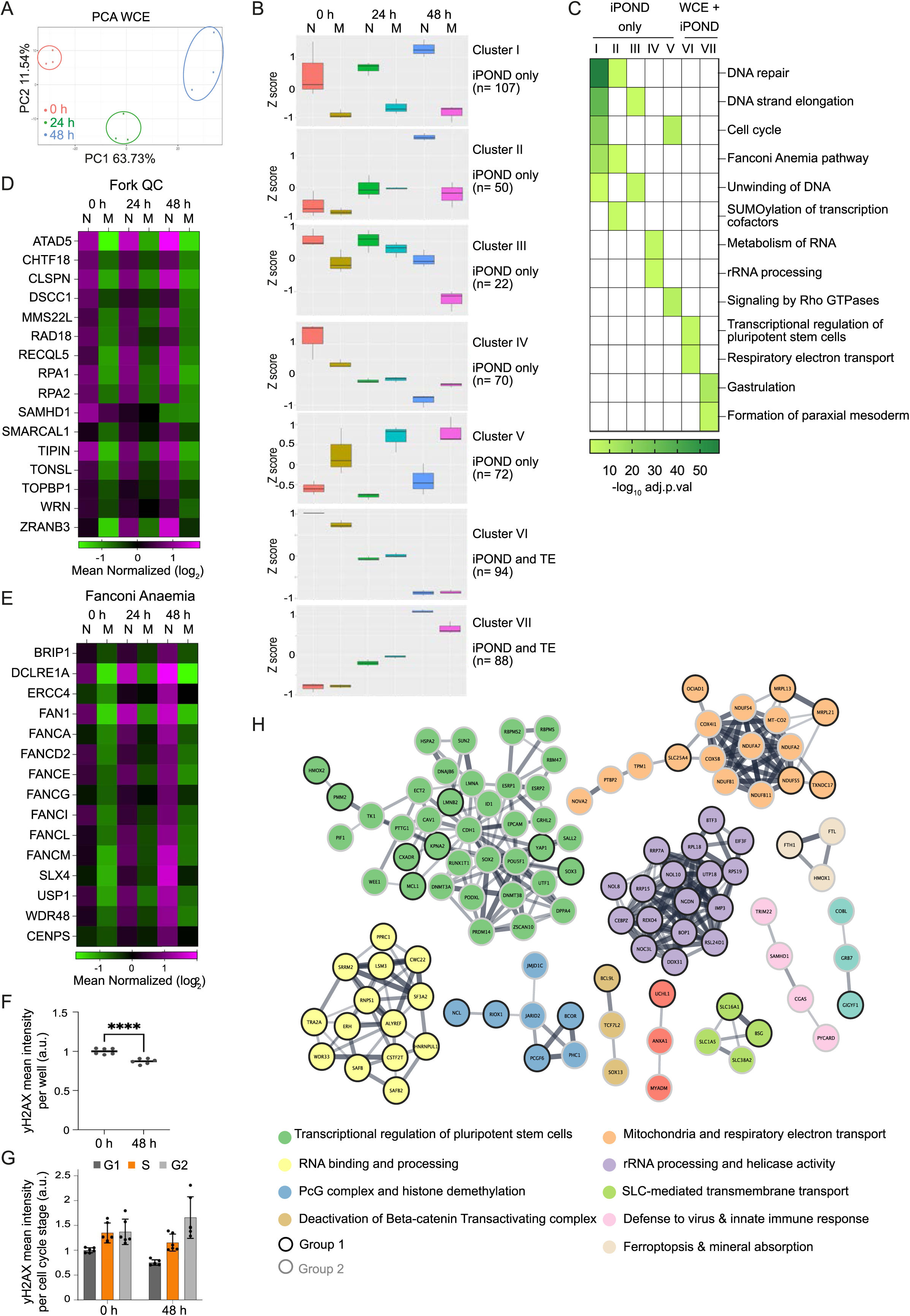
Dynamic changes in the replisome interactome between iPS and PSM cells: TFs are coupled with abundance changes whereas heterochromatin, RNA metabolism and DNA repair are uncoupled. A. Principal component analysis (PCA) of three WCE TMT MS biological replicates based on the proteins identified. PC1 versus PC2 is shown. B. K-Means Clustering of the 503 significantly changing proteins in the iPOND-TMT experiments according to whether they also significantly change in the WCE. Clusters I-V change only in the iPOND experiments whereas clusters VI and VII significantly change in both iPOND and WCE experiments. See also Figure 3C, 3D and Table S4. C. Top Reactome pathway terms (REACC) detected for each cluster from Figure S3B and their representation in all clusters. Significance is shown in increasing green colour reflecting minus log_10_ adjusted p value with 1.3 the minimum significant value and non-significance shown in white. D. Heatmaps showing differential binding of Replication fork quality control factors to nascent and mature chromatin during PSM differentiation. Scale is mean normalized log_2_ (reporter intensities). Protein list is taken from [90]. E. Same as D. but for proteins in the Fanconi anaemia pathway. F. Summary of the relative mean intensity of yH2AX per nucleus per well between iPSCs and PSM 48 h cells. Results from two biological experiments are shown. G. yH2AX level by cell cycle phase relative to the level in G1 phase of iPSCs. EdU positive cells were gated as S phase cells (orange) and cells with low EdU and low DAPI gated as G1 phase (dark grey) and low EdU, high DAPI gated as G2 phase (light grey) as in Figure S1B. Each dot reflects the relative mean nuclear intensity of yH2AX in that cell population. H. STRING interaction network of merged clusters IV and VI, the 164 (70 plus 94) proteins enriched on iPS chromatin. Only high confidence interactions were considered. Network was imported from STRING into Cytoscape and MCL clustering performed with a granularity parameter of 3. Nodes are named and coloured according to the cytoscape clusters. The nodes outlined in black are the proteins from cluster IV that do not change significantly in WCE and only in the iPOND dataset. The nodes outlined in grey are the proteins from cluster VI that change significantly in WCE and in the iPOND data.

**Figure S4.**
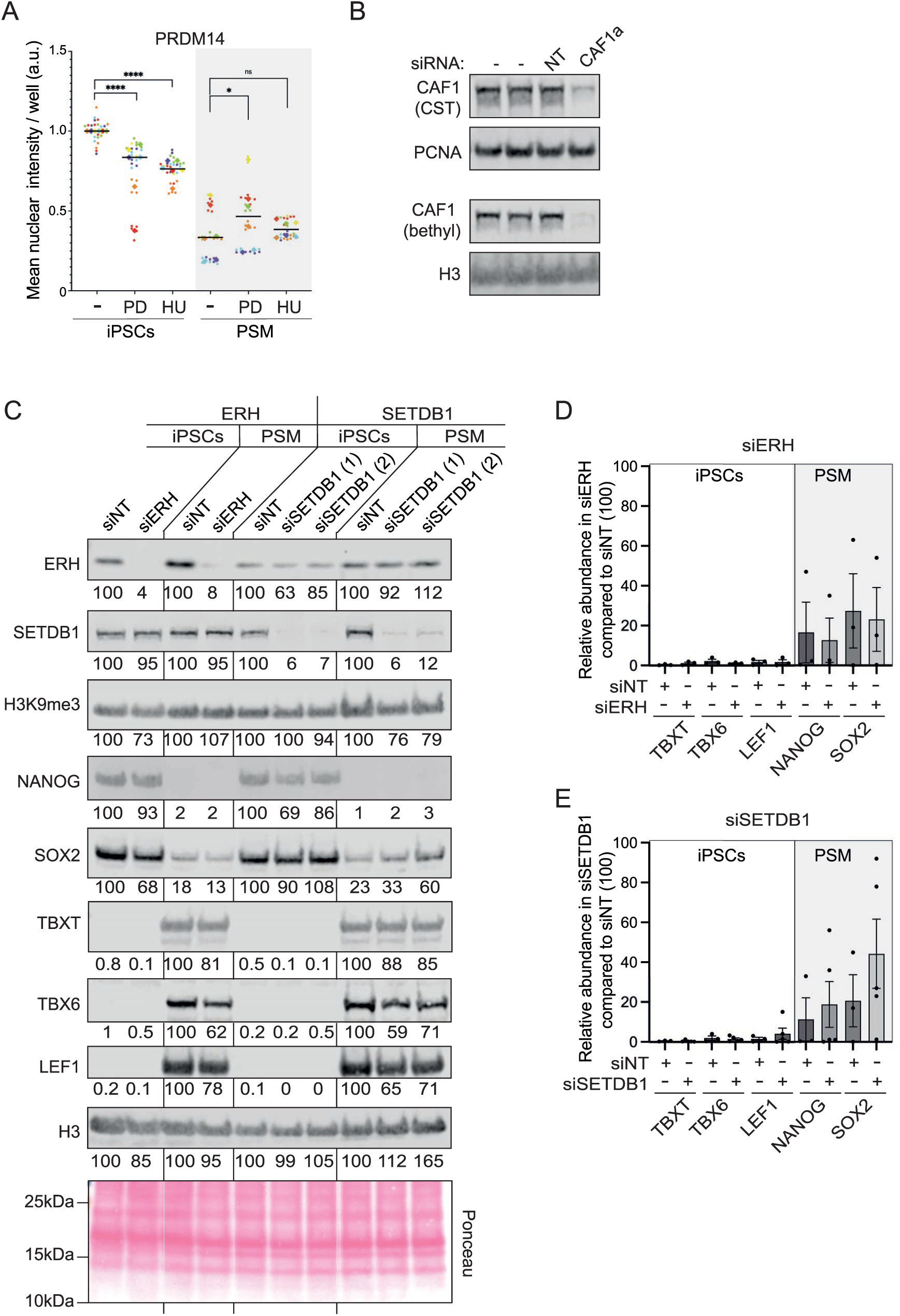
Knockdown of ERH, SETDB1 and CAF1 works efficiently in iPS and PSM cells. A. QIBC analysis showing relative PRDM14 expression in iPSCs and PSM 24-hour cells that are untreated (-) or treated with PD or HU. The results of six biological replicates are shown. B. CAF1 level after siRNA (48 hours) treatment in iPSCs as detected by western blot. C. Example of one representative Western blot experiment on ERH and SETDB1 siRNA samples. D. Summary and quantification of three biological replicates of western blot experiments after ERH siRNA (48 hours differentiation, 72 h siRNA treatment), showing abundance of differentiation markers in iPSCs and pluripotency markers in PSM cells. E. Same as E but for SETDB1 siRNA.

**Table S1.** List of the 503 proteins differentially associated with replicated chromatin during 48 hours of PSM differentiation (iPOND based mass spectrometry). Full dataset available in PRIDE. Project accession number listed in the key resources table.

**Table S2.** List of histone modifications analysed by mass spectrometry in this study. Column A to L, WCE; Column M to AO, iPOND. Full datasets available in PRIDE. Project accession number listed in the key resources table.

**Table S3.** List of 881 proteins significantly changed in abundance within 48 hours of PSM differentiation. (WCE based mass spectrometry). Full dataset available in PRIDE. Project accession number listed in the key resources table.

**Table S4.** Lists of the iPSC and PSM-enriched proteins from groups 1 and 2. Cytoscape cluster name and STRING Description are provided. There are two tabs, one for PSM-enriched proteins (clusters II and VII) and one for iPSC enriched proteins (clusters IV and VI). See also Figures 3C, 3D, S3B and S3H.

## Notes

### Competing Interest Statement

The authors have declared no competing interest.

